# A combination of running and memantine increases neurogenesis and reduces activation of developmentally-born dentate granule neurons in rats

**DOI:** 10.1101/545673

**Authors:** Shaina P. Cahill, Angela Martinovic, John Darby Cole, Jason S. Snyder

**Affiliations:** Department of Psychology, Djavad Mowafaghian Centre for Brain Health, University of British Columbia, 2211 Wesbrook Mall, Vancouver, British Columbia, Canada V6T 2B5

**Keywords:** development, adult neurogenesis, plasticity, immediate early-gene, exercise

## Abstract

During hippocampal-dependent memory formation, sensory signals from the neocortex converge in the dentate gyrus. It is generally believed that the dentate gyrus decorrelates inputs in order to minimize interference between codes for similar experiences, often referred to as pattern separation. The proportion of dentate neurons that are activated by experience is therefore likely to impact how memories are stored and separated. Emerging evidence from mouse models suggests that adult-born neurons can both increase and decrease activity levels in the dentate gyrus. However, the precise conditions that determine the direction of this modulation, and whether it occurs in other species, remains unclear. Furthermore, since the dentate gyrus is composed of a heterogeneous population of cells that are born throughout life, it is unclear if newborn neurons modulate all cells equally. We therefore investigated whether adult neurogenesis in rats modulates activity in dentate gyrus neurons that are born at the peak of early postnatal development. Adult neurogenesis was increased by subjecting rats to an alternating running and memantine treatment schedule, and it was decreased with a transgenic GFAP-TK rat model. Activity was measured by Fos expression in BrdU^+^ cells after rats explored a novel environment. Consistent with an inhibitory role, running+memantine treatment prevented experience-dependent Fos expression in developmentally-born neurons and also in the broader granule cell population. In contrast, blocking neurogenesis did not alter activity patterns. These results suggest that the developmentally-born population of neurons may be a major target of neurogenesis-mediated inhibition. Treatments that promote neurogenesis may therefore benefit disorders where there is elevated activity in the dentate gyrus, such as anxiety and age-related memory impairments.

## 1. Introduction

The dentate gyrus (DG) is the major initial site of convergence of neocortical sensory inputs to the hippocampus [1]. As such, it is uniquely positioned to form associations between the various elements of experience during memory formation. Computational models often emphasize an orthogonalization function of the DG, where similar input patterns are processed and given distinct neural representations, thereby reducing interference between memories [2-4]. Support for this role comes from the unique neuroanatomy of the DG and related circuits, where a relatively large population of DG neurons, with sparse CA3 pyramidal neuron connectivity [5], could allow for distinct ensembles to represent distinct experiences. Also, only a small subset of DG neurons is active in response to a given experience [6-8], and different experiences are represented by distinct DG ensembles [9] and population codes [10]. Factors that regulate the identity and size of the active population of neurons are therefore likely to impact DG functions in memory.

One feature of the DG that sets it apart from other regions of the brain is ongoing neurogenesis throughout adulthood. Adult-born neurons mature through distinct stages where they display enhanced synaptic plasticity [11-13], unique intrinsic electrophysiological profiles [14-16] and maturation and experience-dependent innervation by excitatory and inhibitory inputs [17-21], all of which may contribute to differential recruitment during experience compared to other neurons in the DG [6,17,18]. That adult neurogenesis might regulate DG activity is suggested by structural and physiological evidence that there is a functional balance between cohorts of neurons born at different stages of life, and that adult-born neurons may compete with pre-existing neurons for synaptic inputs [17-19].

Adult-born may also influence DG activity patterns through local excitatory and inhibitory networks. Indeed, inhibition is likely to play a critical role in selecting which DG neurons are to participate in memory encoding [21,22] and a series of studies suggest that adult-born neurons broadly suppress activity in the DG: optogenetic stimulation of adult-born neurons in vitro activates local GABAergic interneurons that inhibit other DG neurons [20,21]. Modulating neurogenesis levels also regulates the degree of activity in the DG network in vitro, such that neurogenesis is associated with lower levels of overall activity [22]. Similar results have been obtained in vivo, where experience-induced recruitment of DG neurons is inversely proportional to neurogenesis levels [20,23,24]. On the other hand, suppressing neurogenesis can also reduce activity in the DG [25] suggesting that, in the intact brain, new neurons might also be capable of driving feedback excitation of the DG.

While neurogenesis regulates DG activity, it remains unclear exactly which neurons are modulated since no study has birthdated the active population of neurons. It is often stated that adult neurogenesis contributes only a small fraction of the total number of DG granule neurons but quantitative estimates suggest the opposite, at least over long intervals [26-28]. Thus, although many DG neurons are born early in life, adult neurogenesis may ultimately contribute a large proportion of the total DG neurons, raising the question of the extent to which developmentally-born neurons are recruited during experience [29]. Indeed, neonatally-born DG neurons undergo apoptosis many months after their birthdate [30], suggesting functional attenuation. Since developmentally-and adult-born neurons have unique cellular properties that are retained even at old cell ages [31], it is important to determine which cohorts of neurons are modulated by neurogenesis in order to understand how the DG, as a whole, contributes to behavior. To address this, we bidirectionally manipulated levels of neurogenesis in rats and examined experience-induced recruitment of neurons that were born at the peak of postnatal development. We additionally examined experience-dependent recruitment of the broader population of DG neurons to determine whether modulatory effects of neurogenesis are specific to the developmentally-born population.

## 2. Methods

### 2.1. Animals and treatments

All procedures were approved by the Animal Care Committee at the University of British Columbia and conducted in accordance with the Canadian Council of Animal Care guidelines regarding humane and ethical treatment of animals. Male experimental Long-Evans rats (Expt. 1) and GFAP-TK transgenic rats (Expt. 2) were bred in the Department of Psychology’s animal facility with a 12-hour light/dark schedule and lights on at 6:00 AM. Different breeders were used for the 2 experiments, with Long Evans rats in Experiment 1 being generated from standard, non-transgenic parents and transgenic GFAP-TK rats in Experiment 2 being generated from a cross between a female GFAP-TK rat and a standard non-transgenic Long Evans male. Breeding occurred in large polyurethane cages (47cm × 37cm × 21cm) containing a polycarbonate tube, aspen chip bedding and ad libitum rat chow and water. The day of birth was designated postnatal day 1. Breeders (both male and female) remained with the litters until P21, when male experimental offspring were weaned into 2 per cage in smaller polyurethane bins (48cm × 27cm × 20cm) with a single polycarbonate tube, aspen chip bedding, and ad libitum rat chow and tap water. In experiment 1 only, animals were weaned into a reverse light-dark cycle (see below).

This study consists of two experiments that investigate how increasing and decreasing adult neurogenesis impact cellular activity in the dentate gyrus, particularly in neurons born during early postnatal development (see timelines in Fig. 1). In both experiments, only male rats were used. They were injected with the thymidine analog BrdU (50 mg/kg, I.P.; Sigma, B500205, St. Louis, MO, USA) on postnatal day 6 (P6) to label neurons born at the peak of granule cell birth [32]. At 2 months of age, neurogenesis manipulation treatments began (see below) and at 6 months of age rats were exposed to a novel context to induce activity-dependent immediate-early gene (IEG) expression (novel context groups). The novel context was an opaque polyurethane cage (47cm × 37cm × 21cm) filled with a mix of aspen chip and corn cob bedding, 2 polycarbonate tubes, 2 paper towels, and 3 ml of white vinegar spread the outer perimeter of the cage. The novel context cage was located in an unfamiliar room. Rats were euthanized 60 minutes after being placed in the novel context. An additional subset of rats did not receive any novel context exposure and was perfused directly from their home cage (home cage groups).

**Figure 1:**
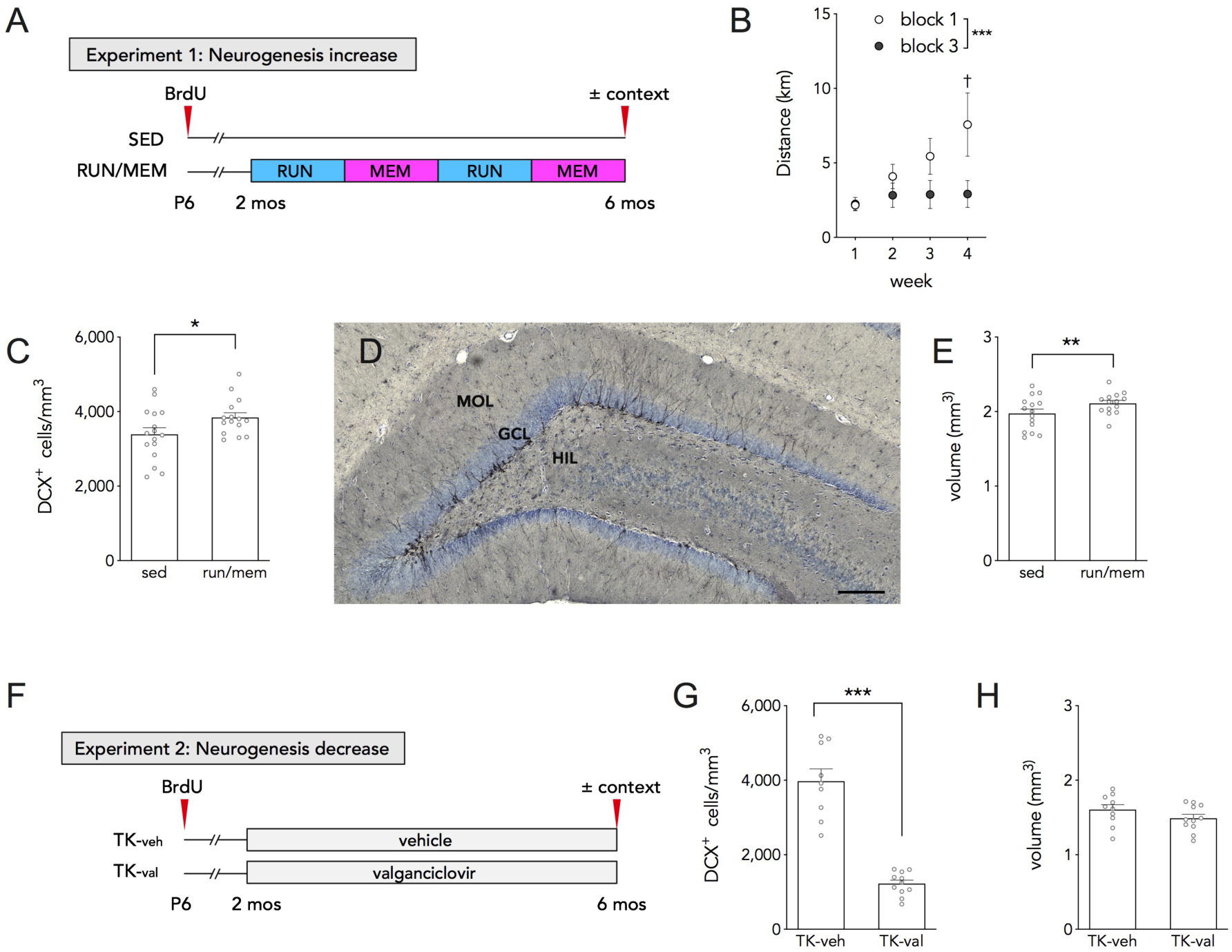
Manipulating adult neurogenesis levels. A) Experiment 1 design and timeline. B) Weekly running distances in blocks 1 and 3. C) More immature, doublecortin^+^ cells (DCX) in RUN/MEM rats than in sedentary controls. D) Representative DCX immunohistochemistry (from a RUN/MEM rat). Scale bar 200 µm. E) RUN/MEM rats had a greater granule cell layer volume than sedentary controls. F) Experiment 2 design and timeline. G) Fewer immature, DCX ^+^ cells in TK-val rats than TK-veh rats. H) TK-val rats had a smaller granule cell layer than TK-veh controls. *P<0.05, **P<0.01, ***P<0.001, ^†^P<0.001 vs block 1 week 1 and all weeks of block 2.

Experiment 1 investigated whether *increasing* adult neurogenesis impacts IEG expression in the DG and specifically in developmentally-born neurons (i.e. born on P6). Neurogenesis was increased by subjecting rats to alternating blocks of intermittent running and memantine injections, as we have done previously [33]. Running + memantine treatment has the greatest neurogenic effect in male rats and, accordingly, only male rats were used in this study. Rats ran during the dark phase because they are more active at this time [34], and because this paradigm matches that of our previous work [33]. Briefly, running wheel cages consisted of a 58cm × 46cm × 38cm plastic tub containing aspen chip bedding, ad libitum rat chow, water, and a 30cm diameter running wheel (Wodent Wheel, Exotic Nutrition). Running distance was measured by tracking wheel revolutions with magnets and a bicycle odometer positioned outside of the cage. Treatment began at 2 months of age, where rats were given 4 x 4-week blocks of treatment (RUN/MEM/RUN/MEM). MEM treatment blocks (2 & 4) consisted of 4 weekly MEM injections (35 mg/kg each, I.P.) and RUN blocks (1 & 3) consisted of rats being placed individually in running wheel cages for 4 hours on weekdays. For the other 2 days per week, animals were pair housed in the running wheel cage with their cage mate, with free access to the running wheels. The timing of RUN treatment was counterbalanced across the 2 rats in a cage, so that each rat would run for the first four hours of the dark phase on one day and the middle four hours of the dark phase on the next day. Sedentary controls were housed in the same colony room as RUN/MEM rats but they remained in their home cages throughout the 4-month treatment period. Sedentary and RUN/MEM rats were exposed to the novel context 1 week after the RUN/MEM rats received their last MEM injection. There was a total of 15 sedentary rats and 16 RUN/MEM rats.

Experiment 2 investigated whether *decreasing* adult neurogenesis impacts IEG expression in the DG and in developmentally-born neurons. Here we used transgenic Long-Evans GFAP-TK (TK) rats, where neurogenesis can be selectively inhibited in adulthood by treatment with the antiviral drug valganciclovir [13,14]. At 2 months of age, some TK rats (n=11) were orally administered 4 mg of valganciclovir (val) to reduce neurogenesis (Hoffman La-Roche; delivered in 0.5 g peanut butter + chow pellets) twice per week for 12 weeks. Rats were not given valganciclovir for the last 4 weeks since we have found that neurogenesis remains suppressed even after ending treatment. Control TK rats (n=10) received 0.5 g vehicle (veh) pellets that did not contain valganciclovir. We refer to neurogenesis-deficient TK rats as the TK-val group and the intact rats as the TK-veh group.

### 2.2. Tissue processing and immunohistochemistry

Immediately following novel context exposure rats were deeply anesthetized with isoflurane and perfused with 4% paraformaldehyde in phosphate buffered saline (PBS, pH 7.4) Brains remained in paraformaldehyde for 48 hr and were then stored in 0.1% sodium azide in PBS until processed. Before processing brains were immersed in 10% glycerol solution for 1 day, 20% glycerol solution for 2 days, and then sectioned coronally at 40 µm on a freezing microtome. Sections were stored in cryoprotectant at - 20°C until immunohistochemical processing. To detect BrdU^+^ and Fos^+^ cells, fluorescent immunohistochemistry was performed on 2 free-floating dorsal DG sections (sections spaced 480 µm apart). Sections were treated in 2N HCl for 30 minutes, incubated at 4^°^C for 3 days in PBS with 10% triton-x, 3% horse serum and goat anti-c-fos primary antibody (1:250; Santa Cruz, sc-52G, Dallas, TX, USA). Sections were then incubated in biotinylated donkey anti-goat secondary antibody (1:250, Thermofisher, A21432, USA) for 60 minutes at room temperature, 5% TSA blocking reagent (Perkin-Elmer, FP1020, Waltmam, MA, USA), Streptavidin-HRP (Perkin-Elmer, NEL750001EA, Waltmam, MA, USA), and then NHS-rhodamine (1:2000, Fisher, PI-46406) in PBS with hydrogen peroxide (1:20,000). Sections were incubated for 3 days in PBS with 10% triton-x, 3% horse serum and mouse anti-BrdU primary antibody (1:200, BD Biosciences; 347580, San Jose, CA, USA). Visualization of BrdU^+^ cells was performed with Alexa 488-conjugated donkey anti-mouse secondary antibody (1:250, Invitrogen/Thermofisher, A21202, USA). Sections were counterstained with DAPI, mounted onto slides and coverslipped with PVA-DABCO.

For DCX analyses four sections, two dorsal and two ventral, were stained for the doublecortin (DCX) to detect immature neurons. Sections were mounted on slides, heated to 90°C in citric acid (0.1M, pH 6.0), washed, permeabilized in PBS with 10% triton-x for 30 min and incubated for three days at 4 °C with goat anti-DCX antibody (1:250 in 10% triton-x and 3% horse serum (sc-8066; Santa Cruz Biotechnology, USA). Sections were washed and incubated in biotinylated donkey anti-goat secondary antibody for 60 minutes (1:250, Jackson, 705065147, West Grove, PA). Cells were then visualized with an avidin-biotin-horseradish peroxidase kit (Vector Laboratories, cat #OK-6100) and cobalt-enhanced 3,3’-diaminobenzidine (DAB) (Sigma Fast Tablets, cat #DO426). Sections were then rinsed in PBS, dehydrated, cleared with citrisolv (Fisher, cat #22143975) and coverslipped with Permount (Fisher, cat #SP15500).

### 2.3. Microscopy and sampling

To evaluate the effectiveness of our neurogenic manipulations, we quantified the number of DG granule cells expressing the immature neuronal marker, DCX [35]. We quantified all DCX^+^ cells across the entire granule cell layer and subgranular zone (∼20 µm wide) from 2 dorsal and 2 ventral sections (8 hemispheres) using a bright field Olympus CX41 microscope and a 40× objective. The granule cell layer volume was calculated by multiplying the section thickness (40 µm) by the 2D area (measured using Stereoinvestigator, MBF systems, Vermont, USA), which was then used to calculate DCX^+^ cell densities.

P6-born BrdU^+^ cells were examined on a confocal microscope (Leica SP8). Expression of the IEG Fos was examined in approximately 200 BrdU^+^ cells per animal, sampled from the suprapyramidal blade of the dorsal dentate gyrus of 2 sections, using a 40x oil-immersion lens (NA 1.3) and 1.5 µm z-sections throughout each tissue section. Image stacks were then analyzed offline. As Fos staining intensity is graded, fluorescence intensity for each cell was measured and compared to background levels within each field (as we have done previously [30]). Fos^+^ cells were marked positive if staining intensity was three times background, which objectively captured cells that had moderate, unambiguous levels of Fos immunostaining as detectable by eye.

To analyze overall IEG expression in the entire granule cell population, approximately 350 DAPI^+^ cells were selected from the same image stacks as the BrdU analyses. Three equal-sized, box-shaped ROIs were spaced along the medio-lateral axis, spanning the full thickness of the granule cell layer (to ensure cells of all ages were sampled equally, as done in [30]). As with BrdU^+^ cells, the Fos signal was measured in each DAPI^+^ cell and counted as Fos^+^ if 3x background. This analysis was performed in 2 different Z-planes in the image stack (each plane separated by 7.5 µm).

Total granule cell layer volume was measured from a 1 in 12 series of cresyl violet-stained sections, using Stereoinvestigator.

### 2.4. Running distance calculations

Running distances were calculated as we have done previously, with individual distances tracked on weekdays during 4-hour blocks [33]. On weekends, when rats were pair housed with the running wheels, individual distances were estimated by assuming the relative distances 2 cage mates ran during the week held for the weekend as well. Weekday and weekend distances were then summed to obtain weekly running distances per rat.

### 2.5. Statistical analyses

Two-way ANOVAs were used to determine whether neurogenic manipulations and behavioral experience altered Fos expression rates in DG neuron populations. Two-way repeated measures ANOVA was used to analyze weekly running behavior across blocks. A one-way ANOVA was performed to analyze treatment effects on granule cell layer volumes. Where significant main effects or interactions were observed, post hoc comparisons were performed using Tukey’s test. For all tests, significance was set at p=0.05.

## 3. Results

### 3.1. Manipulations alter adult neurogenesis and dentate gyrus volume

We quantified DCX^+^ cells to evaluate the long-term efficacy of our manipulations that were intended to increase (Expt 1) and decrease (Expt 2) adult neurogenesis. DCX is expressed for several weeks in young post-mitotic neurons [35] and therefore provides a broad assessment of neurogenesis levels in the weeks prior to examination. In Experiment 1, rats were subjected to alternating RUN and MEM treatments to increase adult neurogenesis. Rats ran significantly more during block 1 than block 3, which was primarily due to a progressive increase in running distances over the course of block 1 (Fig. 1B; repeated measures ANOVA, effect of block F_1, 15_=24.6, P=0.0002; effect of week F_3, 45_=4.6, P=0.007; interaction F_3, 45_=5.8, P=0.0019; Tukey’s comparison of block 1 week 4 vs: block 1 week 1, and block 3 weeks 1-4 all P<0.001).

The RUN/MEM treatment group displayed a 17% increase in DCX^+^ cells compared to controls (Fig. 1C; T_29_=2.065, P=0.048). Whereas we have previously reported that alternating blocks of RUN/MEM increase neurogenesis (DCX^+^ cells) at 2 months in male rats [33], these results indicate that RUN/MEM treatments can result in sustained increases in neurogenesis that are maintained even after 4 months of treatment.

Physical exercise increases hippocampal volume in humans [36]. We therefore examined whether RUN/MEM treatment altered the volume of the granule cell layer. Indeed, we observed a modest but significant 7% increase in granule cell layer volume in RUN/MEM rats compared to sedentary controls (Fig. 1E; T_27_=1.9, P=0.03).

In Experiment 2, TK-val rats were treated with valganciclovir, resulting in a 68% decrease in DCX+ cells compared to TK-veh rats (Fig. 1G; T_18_=8.8, P<0.001). The granule cell layer volume of TK-val rats was reduced by 7% compared to TK-veh rats but this was not statistically significant (Fig. 1H; T_19_=1.4, P=0.09). Of note, the neurogenesis reduction in TK-val rats was less than we have reported [37,38], likely because rats treatment ended 1 month prior to the end of the experiment and neurogenesis may have partially recovered. In addition to differences in volumetric methods (granule cell layer vs whole DG), this may also explain why volume changes were also less robust compared to another TK rats study that inhibited neurogenesis for 4 months [39].

### 3.2. Increasing adult neurogenesis reduces experience-dependent Fos expression in the DG

IEGs are often used as a proxy for neuronal activity since expression profiles match known patterns of electrophysiological activity [40,41], IEGs are expressed in behaviorally-relevant neuronal ensembles [42], and they are required for long-term plasticity and memory [43-45]. To identify whether manipulating adult neurogenesis alters activity in the DG, we quantified Fos expression in P6-born BrdU^+^ cells and DAPI^+^ cells to clarify whether specific populations of DG neurons are recruited after manipulating adult neurogenesis (Fig. 2).

**Figure 2:**
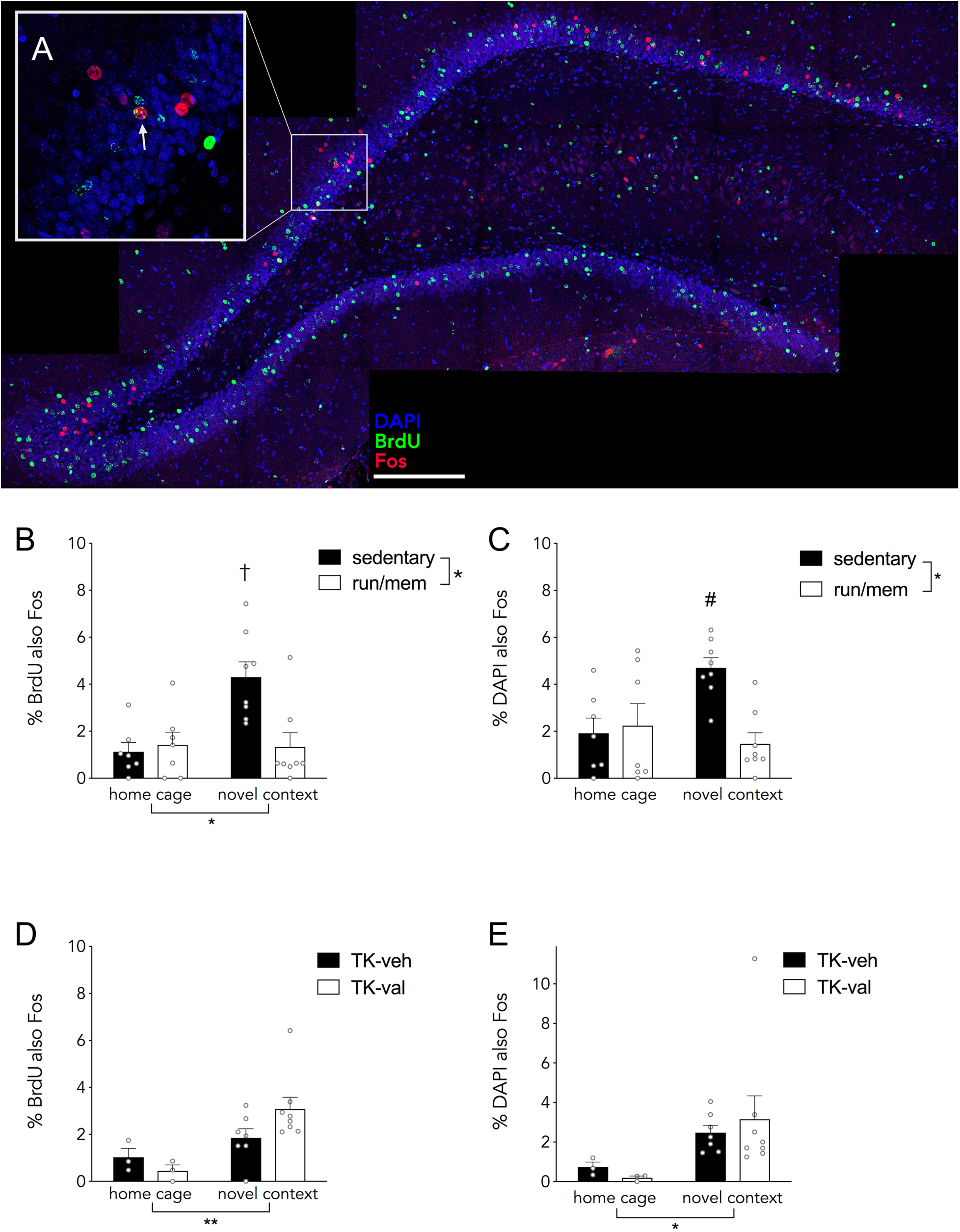
Adult neurogenesis regulates Fos expression in the DG. A) Representative maximum intensity projection image of BrdU/Fos immunostaining in a 40 µm thick section (from a TK-veh rat). Inset shows a BrdU^+^Fos^+^ cell (arrow) from a portion of the z-stack. Increasing neurogenesis with RUN/MEM treatment increases experience-dependent Fos expression in P6-born BrdU^+^ DG neurons (B) and the general DAPI^+^ granule cell population (C). Reducing adult neurogenesis in TK-val rats did not alter experience-dependent Fos expression in P6-born BrdU^+^ DG neurons (D) or the general DAPI^+^ granule cell population (E). *P<0.05, **P<0.01; ^#^P<0.05, ^†^P<0.01 vs RUN/MEM context and sedentary home cage groups.

In Experiment 1, where we increased neurogenesis, there was a main effect of context. Novel context exposure increased the proportion of P6-born BrdU^+^ cells that expressed Fos (Fig. 2B, two-way ANOVA, effect of context F_1, 26_= 7.4, P=0.011). However, this was solely due to an increase in Fos expression in the sedentary, novel-context-exposed rats since Fos expression in novel context-exposed RUN/MEM rats did not differ from home cage controls (effect of treatment F_1, 26_= 5.6, P=0.026; context x treatment interaction F_1, 26_= 8.3, P=0.008; sedentary home cage vs novel context P=0.0028; RUN/MEM home cage vs novel context P=0.99, sedentary novel context vs. RUN/MEM novel context, P=0.0038).

A similar pattern was observed in broader population of DAPI^+^ granule neurons. Animals in the sedentary group showed a significant increase in DAPI^+^Fos^+^ cells after novel context exposure while the RUN/MEM animals failed to show an increase (Fig. 2C; two-way ANOVA context x treatment interaction F_1, 26_= 8.0, P=0.009; sedentary home cage vs novel context P=0.021; RUN/MEM home cage vs novel context P=0.82, sedentary vs. RUN/MEM rats exposed to the novel context, P=0.0046). These results collectively indicate that increasing adult neurogenesis severely blunts activity-dependent activation of the DG and developmentally-born neurons.

### 3.3 Decreasing adult neurogenesis did not alter experience-dependent Fos expression in the DG

To test whether long-term reductions in adult neurogenesis alter DG activity, TK-val rats (reduced neurogenesis) were compared to TK-veh controls (intact neurogenesis). Here, developmentally-born (P6) BrdU^+^ cells displayed increased Fos expression following novel context exposure compared to home cage controls (Fig. 2D; two-way ANOVA, effect of context F_1, 17_= 10.11, P=0.006). We did not observe a significant treatment x context interaction (F_1,17_=2.7, P=0.12) but followed through with a planned comparison to test our hypothesis that adult-born neurons inhibit developmentally-born neurons. However, while novel context-exposed TK-val rats had approximately 60% greater Fos expression in P6-born BrdU^+^ cells than TK-veh controls, this difference was not statistically significant (P=0.19).

To examine whether blocking neurogenesis altered activity in the broader population of DG granule neurons, Fos expression was examined in DAPI^+^ cells. Here, we again found that novel context exposure increased Fos expression compared to the home cage condition, but there was no difference between TK-val and TK-veh rats (Fig. 2E; two-way ANOVA, effect of context F_1, 17_= 4.8, P=0.04; effect of neurogenesis reduction F_1, 17_=0.004, P=0.95; interaction F_1, 17_=0.3, P=0.6).

## 4. Discussion

Here we investigated functional relationships between DG neurons born in adulthood and the peak of DG development. Using RUN/MEM treatment to increase adult neurogenesis [33], and GFAP-TK rats to decrease adult neurogenesis [37], we tested the hypothesis that neurogenesis regulates activity in developmentally-born DG neurons. Indeed, combined RUN/MEM treatment increased immature DCX^+^ cell density and reduced activity in both developmentally-born neurons and the broader granule cell population. The neurogenic efficacy of RUN/MEM treatment at 4 months extends our recent finding that 2 months of treatment increases DCX^+^ cell density in male rats [33]. Thus, it would appear that RUN/MEM is capable of promoting neurogenesis over extended intervals. While this interpretation would be strengthened by demonstrating long-term survival of cells born at earlier stages of treatment, it is bolstered by our finding that the volume of the granule cell layer was greater in RUN/MEM rats than in sedentary controls.

Our data suggest that adult-born neurons reduce activity in populations of DG granule neurons. An alternative explanation is that RUN/MEM rats displayed a lack of experience-induced activity due to direct blockade of NMDA receptors. We feel this is unlikely because MEM treatment was delivered only once per week and an entire week was allowed for washout between the final injection and context exposure. Also, MEM did not alter baseline levels of Fos expression in cage control animals. Finally, others have validated the neurogenic effects of MEM on memory with alternative methods for increasing neurogenesis [46]. Nonetheless, definitive evidence that neurogenesis mediates the effects of RUN/MEM on DG activity could be tested with neurogenesis-deficient animals. Additionally, other methods for increasing neurogenesis could be used to provide converging evidence. For example, future studies could examine whether long term interval running, which selectively promotes neurogenesis in females [33], has greater effects on DG activity in females than in males.

In contrast to the effects of RUN/MEM, there was no significant effect of blocking neurogenesis on DG activity, though there was a trend for more activity in neurogenesis-deficient rats. Little is known about exactly how or when neurogenesis might regulate activity in the DG (discussed below), but the lack of effect in TK-val rats could reflect a floor effect. At 8 months of age, DCX^+^ cell densities in TK-veh rats were less than 10% of what we recently observed in 3 month-old rats [33]. Thus, neurogenesis may have already been too low to modulate DG activity in TK-veh rats, rendering further reductions in TK-val rats insignificant. This would be consistent with findings that 7-month-old mice are behaviorally comparable to irradiated mice in a neurogenesis-dependent task [47], and reports that blocking neurogenesis has greater effects on behavior in younger rodents [48,49].

Since previous studies have examined neurogenesis-mediated modulation of activity in cells of unknown age, in mice, our findings suggest that neurogenesis also modulates DG activity in rats and that adult-born neurons specifically reduce activity in the developmentally-born cohort of neurons. That similar activity profiles were observed in both P6-born DG neurons and DAPI^+^ cells provides novel evidence that the global pattern of activity in the DG is representative of the neonatally-born population, though further experiments are needed to determine whether all DG granule neurons, born at various stages of life, display similar experience-dependent activity patterns.

### 4.1. Adult neurogenesis and regulation of activity in the hippocampus

Our findings raise questions about how and when neurogenesis modulates activity in the DG. Currently, little is known about the mechanisms by which newborn neurons interact with other neuronal populations during behavior. However, some possibilities can be entertained based on anatomy, physiology, and patterns of experience-dependent neuronal recruitment.

One mechanism for activity regulation could be through competitive dynamics at afferent synaptic inputs onto DG neurons. Adult-born neurons initially share afferent presynaptic terminals with existing DG neurons [19] and there is a causal balance between existing and new synaptic inputs in the DG: whereas reducing spines on mature neurons promotes integration of immature neurons [17], increasing and decreasing adult neurogenesis induces compensatory changes in synaptic drive onto mature neurons located in the superficial granule cell layer [18]. Assuming a competitive model, a large number of new neurons in RUN/MEM rats could have outcompeted developmentally-born neurons for synaptic inputs, thereby reducing experience-dependent Fos expression. However, this account is unlikely to fully explain our findings since increased numbers of synaptically-integrated new neurons would have presumably been reflected by elevated Fos expression in DAPI^+^ cells.

Adult neurogenesis may also modulate DG activity through local circuitry. For example, experience-dependent neuronal recruitment in the DG is tightly controlled by inhibition [50]. Since adult-born neurons drive local interneurons [20,21,51], it is possible that elevated neurogenesis in the RUN/MEM group could have suppressed DG activity through lateral or feedback inhibition. Similarly, blocking [23] or silencing [24] adult-born neurons in mice may increase DG activity through a disinhibition. While lateral or feedback excitation circuits in the DG are not as common or well understood, they could also contribute to neurogenesis-medicated modulation of DG activity. For example, adult-born neurons could trigger feedback excitation via mossy cells [51,52] or CA3 pyramidal neurons [54,55], which may explain why blocking neurogenesis reduces DG activity in mice that are trained in an unpredictable fear conditioning paradigm [25].

Finally, DG activity could be regulated by certain internal or behavioral states, such as stress or fear. Burghardt et al. found that irradiated mice had elevated DG Arc expression when trained on an aversive spatial reversal task [23]. However, irradiated mice were also slower to learn the reversal, experienced many more footshocks as a result, and therefore may have experienced greater levels of fear. Conversely, neurogenesis-deficient mice displayed less fear in an ambiguous aversive conditioning paradigm and had lower levels of Fos^+^ cells in the DG [25]. Chemogenetic silencing of adult-born neurons also promotes fearful social behavior and elevates DG Fos levels [24]; while silencing new neurons could have elevated DG activity through direct circuit interactions, which in turn cause fear behavior, it is also possible that silencing new neurons increased fear/reduced social exploration, which then elevated DG activity. By this interpretation, in the current study, RUN/MEM may have reduced fearfulness of the novel environment, either by increasing neurogenesis or due to the repeated injections and handling, which would have subsequently lowered DG activity. Also, similar Fos levels in TK-veh and TK-val rats is consistent with findings that neurogenesis-intact and deficient animals typically display similar levels of fear/anxiety at baseline in a novel environment [53-56].

Our results are relevant for current models that propose cell-age-dependent functions for DG neurogenesis. Specifically, it has been proposed that DG neurons pass through immature stages when they are selectively recruited to encode memories for events that occur during a critical period [3,57], and that DG neurons may become functionally silent with age [29]. That P6-born DG neurons (in sedentary rats and TK rats) upregulated Fos following exposure to a novel environment suggests that old DG neurons are fully capable of encoding new experiences, though this could reflect the rather impoverished environment, and lack of opportunity for learning, that rats experienced for the majority of their lives. It will therefore be important for future experiments to more rigorously examine how cohorts of immature vs mature cells encode experiences that occur at different intervals throughout the life.

### 4.2. Implications for cognition and mental health

Theoretical models predict that sparse DG activity allows for separation of incoming sensory signals into distinct population codes, which serves to reduce memory interference [2-4]. While most behavioral studies have not directly measured DG activity levels, there is a wealth of evidence that DG neurogenesis is particularly important for precise memory under conditions of high interference [58-64]. An inhibitory role for new neurons may therefore be a viable approach to promote sparse coding schemes in the DG and improve the specificity and accuracy of memory in disorders such as generalized fear [65]. Such a role is also relevant for age-related memory decline, since neurogenesis declines precipitously with age [31,66,67] and early age-related deficits in mnemonic discrimination are associated with DG-CA3 hyperactivity [68].

The DG [69] and neurogenesis [53,56] have also been implicated in anxiety. While anxiety is typically considered to be an innate (rather than learned) fear, it also lacks specificity and therefore resembles in some respects the generalized fear that can be observed in discriminative conditioning paradigms. Anxiety is also associated with more widespread patterns of IEG expression in the DG, which can be normalized by exercise-induced increases in GABAergic signalling [70]. Since exercise potently upregulates neurogenesis this suggests that new neurons may be involved, though recent work suggests exercise can be anxiolytic in the absence of neurogenesis [71]. Given the complexity of the neurogenesis-based modulation of DG activity discussed above, more experiments are warranted to determine if neurogenesis modulates anxiety in some behavioral states and not others.

## 5. Conclusions

Previous evidence for neurogenesis-mediated modulation of DG activity comes solely from mouse models, often using transgenic and optogenetic methods that are less amenable to rat models and unlikely to be used in humans. We therefore adopted a neurogenic approach that is more translatable to human conditions, and provide support for a role of neurogenesis in DG inhibition. Neurogenic therapies, such as exercise, MEM or other compounds such as P7C3 (which has recently been validated in nonhuman primates [72]) could therefore be viable approaches for treating cognitive and mental health disorders that are associated with elevated activity rates in the DG.

## Acknowledgements

This research was supported by a Discovery Grant from the Natural Sciences and Engineering Research Council of Canada (NSERC; JSS), a New Investigator Award from the Canadian Institutes of Health Research (JSS), a Scholar award from the Michael Smith Foundation for Health Research (JSS) and Killam and NSERC graduate scholarships (SPC).

## References

[1] D.G. Amaral, H.E. Scharfman, P. Lavenex, The dentate gyrus: fundamental neuroanatomical organization (dentate gyrus for dummies), Prog Brain Res. 163 (2007) 3–22. doi:10.1016/S0079-6123(07)63001-5.

[2] A. Treves, E.T. Rolls, Computational constraints suggest the need for two distinct input systems to the hippocampal CA3 network, Hippocampus. 2 (1992) 189–199. doi:10.1002/hipo.450020209.

[3] J.B. Aimone, J. Wiles, F.H. Gage, Computational influence of adult neurogenesis on memory encoding, Neuron. 61 (2009) 187–202. doi:10.1016/j.neuron.2008.11.026.

[4] J.L. McClelland, N.H. Goddard, Considerations arising from a complementary learning systems perspective on hippocampus and neocortex, Hippocampus. 6 (1996) 654–665. doi:10.1002/(SICI)1098-1063(1996)6:6<654::AID-HIPO8>3.0.CO;2-G.

[5] L. Acsády, A. Kamondi, A. Sík, T. Freund, G. Buzsáki, GABAergic cells are the major postsynaptic targets of mossy fibers in the rat hippocampus, J Neurosci. 18 (1998) 3386–3403.

[6] J.S. Snyder, R. Radik, J.M. Wojtowicz, H.A. Cameron, Anatomical gradients of adult neurogenesis and activity: young neurons in the ventral dentate gyrus are activated by water maze training, Hippocampus. 19 (2009) 360–370. doi:10.1002/hipo.20525.

[7] A. Thome, D.F. Marrone, T.M. Ellmore, M.K. Chawla, P. Lipa, V. Ramirez-Amaya, et al., Evidence for an Evolutionarily Conserved Memory Coding Scheme in the Mammalian Hippocampus, J Neurosci. (2017) 3057–16. doi:10.1523/JNEUROSCI.3057-16.2017.

[8] D. GoodSmith, X. Chen, C. Wang, S.H. Kim, H. Song, A. Burgalossi, et al., Spatial Representations of Granule Cells and Mossy Cells of the Dentate Gyrus, Neuron. 93 (2017) 677–690.e5. doi:10.1016/j.neuron.2016.12.026.

[9] E. satvat, B. Schmidt, M. Argraves, D.F. Marrone, E.J. Markus, Changes in task demands alter the pattern of zif268 expression in the dentate gyrus, J Neurosci. 31 (2011) 7163–7167. doi:10.1523/JNEUROSCI.0094-11.2011.

[10] J.P. Neunuebel, J.J. Knierim, CA3 Retrieves Coherent Representations from Degraded Input: Direct Evidence for CA3 Pattern Completion and Dentate Gyrus Pattern Separation, Neuron. 81 (2014) 416–427. doi:10.1016/j.neuron.2013.11.017.

[11] J.S. Snyder, N. Kee, J.M. Wojtowicz, Effects of adult neurogenesis on synaptic plasticity in the rat dentate gyrus, J Neurophysiol. 85 (2001) 2423–2431.

[12] S. Ge, C.-H. Yang, K.-S. Hsu, G.-L. Ming, H. Song, A Critical Period for Enhanced Synaptic Plasticity in Newly Generated Neurons of the Adult Brain, Neuron. 54 (2007) 559–566. doi:10.1016/j.neuron.2007.05.002.

[13] C. Schmidt-Hieber, P. Jonas, J. Bischofberger, Enhanced synaptic plasticity in newly generated granule cells of the adult hippocampus, Nature. 429 (2004) 184–187. doi:10.1038/nature02553.

[14] M.S. Espósito, V.C. Piatti, D.A. Laplagne, N.A. Morgenstern, C.C. Ferrari, F.J. Pitossi, et al., Neuronal differentiation in the adult hippocampus recapitulates embryonic development, J Neurosci. 25 (2005) 10074–10086. doi:10.1523/JNEUROSCI.3114-05.2005.

[15] J. Brunner, M. Neubrandt, S. Van-Weert, T. Andrási, F.B. Kleine Borgmann, S. Jessberger, et al., Adult-born granule cells mature through two functionally distinct states, eLife. 3 (2014) e03104. doi:10.7554/eLife.03104.

[16] L.A. Mongiat, M.S. Espósito, G. Lombardi, A.F. Schinder, Reliable activation of immature neurons in the adult hippocampus, PLoS ONE. 4 (2009) e5320. doi:10.1371/journal.pone.0005320.

[17] K.M. McAvoy, K.N. Scobie, S. Berger, C. Russo, N. Guo, P. Decharatanachart, et al., Modulating Neuronal Competition Dynamics in the Dentate Gyrus to Rejuvenate Aging Memory Circuits, Neuron. 91 (2016) 1356–1373. doi:10.1016/j.neuron.2016.08.009.

[18] E.W. Adlaf, R.J. Vaden, A.J. Niver, A.F. Manuel, V.C. Onyilo, M.T. Araujo, et al., Adult-born neurons modify excitatory synaptic transmission to existing neurons, eLife. 6 (2017) e19886. doi:10.7554/eLife.19886.

[19] N. Toni, E.M. Teng, E.A. Bushong, J.B. Aimone, C. Zhao, A. Consiglio, et al., Synapse formation on neurons born in the adult hippocampus, Nat Neurosci. 10 (2007) 727–734. doi:10.1038/nn1908.

[20] L.J. Drew, M.A. Kheirbek, V.M. Luna, C.A. Denny, M.A. Cloidt, M.V. Wu, et al., Activation of local inhibitory circuits in the dentate gyrus by adult-born neurons, Hippocampus. (2015). doi:10.1002/hipo.22557.

[21] S.G. Temprana, L.A. Mongiat, S.M. Yang, M.F. Trinchero, D.D. Alvarez, E. Kropff, et al., Delayed Coupling to Feedback Inhibition during a Critical Period for the Integration of Adult-Born Granule Cells, Neuron. 85 (2015) 116–130. doi:10.1016/j.neuron.2014.11.023.

[22] T. Ikrar, N. Guo, K. He, A. Besnard, S. Levinson, A. Hill, et al., Adult neurogenesis modifies excitability of the dentate gyrus, Front Neural Circuits. 7 (2013) 204. doi:10.3389/fncir.2013.00204.

[23] N.S. Burghardt, E.H. Park, R. Hen, A.A. Fenton, Adult-born hippocampal neurons promote cognitive flexibility in mice, Hippocampus. 22 (2012) 1795–1808. doi:10.1002/hipo.22013.

[24] C. Anacker, V.M. Luna, G.S. Stevens, A. Millette, R. Shores, J.C. Jimenez, et al., Hippocampal neurogenesis confers stress resilience by inhibiting the ventral dentate gyrus, Nature. 559 (2018) 1–22. doi:10.1038/s41586-018-0262-4.

[25] L.R. Glover, T.J. Schoenfeld, R.-M. Karlsson, D.M. Bannerman, H.A. Cameron, Ongoing neurogenesis in the adult dentate gyrus mediates behavioral responses to ambiguous threat cues, PLoS Biol. 15 (2017) e2001154. doi:10.1371/journal.pbio.2001154.

[26] J.S. Snyder, H.A. Cameron, Could adult hippocampal neurogenesis be relevant for human behavior? Behav Brain Res. 227 (2012) 384–390. doi:10.1016/j.bbr.2011.06.024.

[27] K.L. Spalding, O. Bergmann, K. Alkass, S. Bernard, M. Salehpour, H.B. Huttner, et al., Dynamics of hippocampal neurogenesis in adult humans, Cell. 153 (2013) 1219–1227. doi:10.1016/j.cell.2013.05.002.

[28] N.A. DeCarolis, M. Mechanic, D. Petrik, A. Carlton, J.L. Ables, S. Malhotra, et al., In vivo contribution of nestin- and GLAST-lineage cells to adult hippocampal neurogenesis, Hippocampus. 23 (2013) 708–719. doi:10.1002/hipo.22130.

[29] C.B. Alme, R.A. Buzzetti, D.F. Marrone, J.K. Leutgeb, M.K. Chawla, M.J. Schaner, et al., Hippocampal granule cells opt for early retirement, Hippocampus. 20 (2010) 1109–1123. doi:10.1002/hipo.20810.

[30] S.P. Cahill, R.Q. Yu, D. Green, E.V. Todorova, J.S. Snyder, Early survival and delayed death of developmentally-born dentate gyrus neurons, Hippocampus. (2017). doi:10.1002/hipo.22760.

[31] J.S. Snyder, Recalibrating the Relevance of Adult Neurogenesis, Trends Neurosci. (2019) 1–15. doi:10.1016/j.tins.2018.12.001.

[32] A.R. Schlessinger, W.M. Cowan, D.I. Gottlieb, An autoradiographic study of the time of origin and the pattern of granule cell migration in the dentate gyrus of the rat, J Comp Neurol. 159 (1975) 149–175. doi:10.1002/cne.901590202.

[33] S.P. Cahill, J.D. Cole, R.Q. Yu, J. Clemans-Gibbon, J.S. Snyder, Differential Effects of Extended Exercise and Memantine Treatment on Adult Neurogenesis in Male and Female Rats, Neuroscience. 390 (2018) 241–255. doi:10.1016/j.neuroscience.2018.08.028.

[34] M.M. Holmes, L.A.M. Galea, R.E. Mistlberger, G. Kempermann, Adult hippocampal neurogenesis and voluntary running activity: circadian and dose-dependent effects, J Neurosci Res. 76 (2004) 216–222. doi:10.1002/jnr.20039.

[35] J.P. Brown, S. Couillard-Després, C.M. Cooper-Kuhn, J. Winkler, L. Aigner, H.G. Kuhn, Transient expression of doublecortin during adult neurogenesis, J Comp Neurol. 467 (2003) 1–10. doi:10.1002/cne.10874.

[36] K.I. Erickson, M.W. Voss, R.S. Prakash, C. Basak, A. Szabo, L. Chaddock, et al., Exercise training increases size of hippocampus and improves memory, Proceedings of the National Academy of Sciences. 108 (2011) 3017–3022. doi:10.1073/pnas.1015950108.

[37] J.S. Snyder, L. Grigereit, A. Russo, D.R. Seib, M. Brewer, J. Pickel, et al., A Transgenic Rat for Specifically Inhibiting Adult Neurogenesis, eNeuro. 3 (2016). doi:10.1523/ENEURO.0064-16.2016.

[38] D.R. Seib, E. Chahley, O. Princz-Lebel, J.S. Snyder, Intact memory for local and distal cues in male and female rats that lack adult neurogenesis, PLoS ONE. 13 (2018) e0197869–15. doi:10.1371/journal.pone.0197869.

[39] T.J. Schoenfeld, H.C. McCausland, H.D. Morris, V. Padmanaban, H.A. Cameron, Stress and loss of adult neurogenesis differentially reduce hippocampal volume, Biol Psychiatry. 82 (2017) 1–34. doi:10.1016/j.biopsych.2017.05.013.

[40] M.K. Chawla, J.F. Guzowski, V. Ramirez-Amaya, P. Lipa, K.L. Hoffman, L.K. Marriott, et al., Sparse, environmentally selective expression of Arc RNA in the upper blade of the rodent fascia dentata by brief spatial experience, Hippocampus. 15 (2005) 579–586. doi:10.1002/hipo.20091.

[41] J.F. Guzowski, B.L. McNaughton, C.A. Barnes, P.F. Worley, Environment-specific expression of the immediate-early gene Arc in hippocampal neuronal ensembles, Nat Neurosci. 2 (1999) 1120–1124. doi:10.1038/16046.

[42] X. Liu, S. Ramirez, P.T. Pang, C.B. Puryear, A. Govindarajan, K. Deisseroth, et al., Optogenetic stimulation of a hippocampal engram activates fear memory recall, Nature. 484 (2012) 381–385. doi:10.1038/nature11028.

[43] M.W. Jones, M.L. Errington, P.J. French, A. Fine, T.V. Bliss, S. Garel, et al., A requirement for the immediate early gene Zif268 in the expression of late LTP and long-term memories, Nat Neurosci. 4 (2001) 289–296. doi:10.1038/85138.

[44] R.A. Countryman, N.L. Kaban, P.J. Colombo, Hippocampal c-fos is necessary for long-term memory of a socially transmitted food preference, Neurobiol Learn Mem. 84 (2005) 175–183. doi:10.1016/j.nlm.2005.07.005.

[45] J.F. Guzowski, G.L. Lyford, G.D. Stevenson, F.P. Houston, J.L. McGaugh, P.F. Worley, et al., Inhibition of activity-dependent arc protein expression in the rat hippocampus impairs the maintenance of long-term potentiation and the consolidation of long-term memory, J Neurosci. 20 (2000) 3993–4001.

[46] K.G. Akers, A. Martinez-Canabal, L. Restivo, A.P. Yiu, A. De Cristofaro, H.-L. Hsiang, et al., Hippocampal Neurogenesis Regulates Forgetting During Adulthood and Infancy, Science. 344 (2014) 598–602. doi:10.1126/science.1248903.

[47] C.A. Denny, N.S. Burghardt, D.M. Schachter, R. Hen, M.R. Drew, 4-to 6-week-old adult-born hippocampal neurons influence novelty-evoked exploration and contextual fear conditioning, Hippocampus. 22 (2012) 1188–1201. doi:10.1002/hipo.20964.

[48] A. Martinez-Canabal, K.G. Akers, S.A. Josselyn, P.W. Frankland, Age-dependent effects of hippocampal neurogenesis suppression on spatial learning, Hippocampus. 23 (2013) 66–74. doi:10.1002/hipo.22054.

[49] L. Wei, M.J. Meaney, R.S. Duman, A. Kaffman, Affiliative behavior requires juvenile, but not adult neurogenesis, J Neurosci. 31 (2011) 14335–14345. doi:10.1523/JNEUROSCI.1333-11.2011.

[50] T. Stefanelli, C. Bertollini, C. Lüscher, D. Muller, P. Mendez, Hippocampal Somatostatin Interneurons Control the Size of Neuronal Memory Ensembles, Neuron. 89 (2016) 1074–1085. doi:10.1016/j.neuron.2016.01.024.

[51] P.S. Buckmaster, H.J. Wenzel, D.D. Kunkel, P.A. Schwartzkroin, Axon arbors and synaptic connections of hippocampal mossy cells in the rat in vivo, J Comp Neurol. 366 (1996) 271–292.

[52] P.S. Buckmaster, B.W. Strowbridge, D.D. Kunkel, D.L. Schmiege, P.A. Schwartzkroin, Mossy cell axonal projections to the dentate gyrus molecular layer in the rat hippocampal slice, Hippocampus. 2 (1992) 349–362. doi:10.1002/hipo.450020403.

[53] J.S. Snyder, A. Soumier, M. Brewer, J. Pickel, H.A. Cameron, Adult hippocampal neurogenesis buffers stress responses and depressive behaviour, Nature. 476 (2011) 458–461. doi:10.1038/nature10287.

[54] L. Santarelli, M. Saxe, C. Gross, A. Surget, F. Battaglia, S. Dulawa, et al., Requirement of hippocampal neurogenesis for the behavioral effects of antidepressants, Science. 301 (2003) 805–809. doi:10.1126/science.1083328.

[55] A. Surget, M. Saxe, S. Leman, Y. Ibarguen-Vargas, S. Chalon, G. Griebel, et al., Drug-dependent requirement of hippocampal neurogenesis in a model of depression and of antidepressant reversal, Biol Psychiatry. 64 (2008) 293–301. doi:10.1016/j.biopsych.2008.02.022.

[56] J.-M. Revest, D. Dupret, M. Koehl, C. Funk-Reiter, N. Grosjean, P.V. Piazza, et al., Adult hippocampal neurogenesis is involved in anxiety-related behaviors, Mol Psychiatry. 14 (2009) 959–967. doi:10.1038/mp.2009.15.

[57] R. Finnegan, S. Becker, Neurogenesis paradoxically decreases both pattern separation and memory interference, Front. Syst. Neurosci. 9 (2015) 136. doi:10.3389/fnsys.2015.00136.

[58] P. Luu, O.C. Sill, L. Gao, S. Becker, J.M. Wojtowicz, D.M. Smith, The role of adult hippocampal neurogenesis in reducing interference, Behav Neurosci. 126 (2012) 381–391. doi:10.1037/a0028252.

[59] G. Winocur, S. Becker, P. Luu, S. Rosenzweig, J.M. Wojtowicz, Adult hippocampal neurogenesis and memory interference, Behav Brain Res. 227 (2012) 464–469. doi:10.1016/j.bbr.2011.05.032.

[60] C.D. Clelland, M. Choi, C. Romberg, G.D. Clemenson, A. Fragniere, P. Tyers, et al., A functional role for adult hippocampal neurogenesis in spatial pattern separation, 325 (2009) 210–213. doi:10.1126/science.1173215.

[61] A. Sahay, K.N. Scobie, A.S. Hill, C.M. O’Carroll, M.A. Kheirbek, N.S. Burghardt, et al., Increasing adult hippocampal neurogenesis is sufficient to improve pattern separation, Nature. 472 (2011) 466–470. doi:10.1038/nature09817.

[62] M.A. Kheirbek, L. Tannenholz, R. Hen, NR2B-dependent plasticity of adult-born granule cells is necessary for context discrimination, J Neurosci. 32 (2012) 8696–8702. doi:10.1523/JNEUROSCI.1692-12.2012.

[63] J.R. Epp, R. Silva Mera, S. Köhler, S.A. Josselyn, P.W. Frankland, Neurogenesis-mediated forgetting minimizes proactive interference, Nat Comms. 7 (2016) 10838. doi:10.1038/ncomms10838.

[64] S. Tronel, L. Belnoue, N. Grosjean, J.-M. Revest, P.-V. Piazza, M. Koehl, et al., Adult-born neurons are necessary for extended contextual discrimination, Hippocampus. 22 (2012) 292–298. doi:10.1002/hipo.20895.

[65] A. Besnard, A. Sahay, Adult Hippocampal Neurogenesis, Fear Generalization, and Stress, Neuropsychopharmacology. 41 (2016) 24–44. doi:10.1038/npp.2015.167.

[66] J. Gil-Mohapel, P.S. Brocardo, W. Choquette, R. Gothard, J.M. Simpson, B.R. Christie, Hippocampal neurogenesis levels predict WATERMAZE search strategies in the aging brain, PLoS ONE. 8 (2013) e75125. doi:10.1371/journal.pone.0075125.

[67] J. Altman, G.D. Das, Autoradiographic and histological evidence of postnatal hippocampal neurogenesis in rats, J Comp Neurol. 124 (1965) 319–335.

[68] Z.M. Reagh, J.A. Noche, N.J. Tustison, D. Delisle, E.A. Murray, M.A. Yassa, Functional Imbalance of Anterolateral Entorhinal Cortex and Hippocampal Dentate/CA3 Underlies Age-Related Object Pattern Separation Deficits, Neuron. 97 (2018) 1187–1198.e4. doi:10.1016/j.neuron.2018.01.039.

[69] M.A. Kheirbek, L.J. Drew, N.S. Burghardt, D.O. Costantini, L. Tannenholz, S.E. Ahmari, et al., Differential Control of Learning and Anxiety along the Dorsoventral Axis of the Dentate Gyrus, Neuron. 77 (2013) 955–968. doi:10.1016/j.neuron.2012.12.038.

[70] T.J. Schoenfeld, P. Rada, P.R. Pieruzzini, B. Hsueh, E. Gould, Physical exercise prevents stress-induced activation of granule neurons and enhances local inhibitory mechanisms in the dentate gyrus, J Neurosci. 33 (2013) 7770–7777. doi:10.1523/JNEUROSCI.5352-12.2013.

[71] T.J. Schoenfeld, H.C. McCausland, A.N. Sonti, H.A. Cameron, Anxiolytic Actions of Exercise in Absence of New Neurons, Hippocampus. 26 (2016) 1373–1378. doi:10.1002/hipo.22649.

[72] M.D. Bauman, C.M. Schumann, E.L. Carlson, S.L. Taylor, E. Vázquez-Rosa, C.J. Cintrón-Pérez, et al., Neuroprotective efficacy of P7C3 compounds in primate hippocampus, Transl Psychiatry. 8 (2018) 1–11. doi:10.1038/s41398-018-0244-1.

